# DNA-damage induced cell death in *yap1;wwtr1* mutant epidermal basal cells

**DOI:** 10.1101/2021.07.23.453490

**Authors:** Jason K. H. Lai, Pearlyn J. Y. Toh, Hamizah A. Cognart, Geetika Chouhan, Timothy E. Saunders

**Author notes:** Corresponding authors: J. K. H. L., T. E. S.

## Abstract

In a previous study, it was reported that Yap1 and Wwtr1 in zebrafish regulates the morphogenesis of the posterior body and epidermal fin fold (Kimelman, D., et al. 2017). We report here that DNA damage induces apoptosis of epidermal basal cells (EBCs) in zebrafish *yap1^−/−^;wwtr1^−/−^* embryos. Specifically, these mutant EBCs exhibit active Caspase-3, Caspase-8 and γH2AX, consistent with DNA damage serving as a stimulus of the extrinsic apoptotic pathway in epidermal cells. Live imaging of zebrafish epidermal cells reveals a steady growth of basal cell size in the developing embryo, but this growth is inhibited in mutant basal cells followed by apoptosis, leading to the hypothesis that factors underscoring cell size play a role in this DNA damage-induced apoptosis phenotype. We tested two of these factors using cell stretching and substrate stiffness assays, and found that HaCaT cells cultured on stiff substrates exhibit more numerous γH2AX foci compared to ones cultured on soft substrates. Thus, we propose that substrate rigidity modulates genomic stress in the developing epidermal cell, and that Yap1 and Wwtr1 are required for its survival.

## INTRODUCTION

The Hippo signalling pathway is widely known for its role in regulating tissue and organ size (Johnson & Halder, 2014). More recently, the roles of its downstream effectors, Yap1 and Wwtr1 (or widely referred to as Taz), have been described in a variety of developing tissues. In the developing heart, Wwtr1 participates in cardiomyocyte decision-making to become a trabecular cell (Lai et al., 2018). Additionally, in the cardiovascular system, YAP1 and WWTR1 are important for sprouting angiogenesis (Astone et al., 2018; Kim et al., 2017; Neto et al., 2018; Wang et al., 2017) and vascular stability (Nakajima et al., 2017). Yap1 and Wwtr1 also regulate the retinal epithelium cell fate in the eye (Miesfeld et al., 2015), kidney branching morphogenesis (Reginensi et al., 2015) and fibronectin assembly for tail morphogenesis (Kimelman et al., 2017). In this report, we show that Yap1 and Wwtr1 are required for the survival and growth of epidermal cells during skin development.

In the zebrafish, the non-neural ectoderm, marked by *ΔNp63*, emerges from the ventral side of the gastrulating embryo, and spreads dorsally to cover the entire embryo as a sheet of cells beneath the periderm during segmentation stages (Bakkers et al., 2002; Solnica-Krezel, 2005). Residing in this layer are the epidermal basal cells (EBCs) which are responsible for epidermal cell renewal and homeostasis (Blanpain & Fuchs, 2006). In mice, Yap1 and Wwtr1, acting downstream of integrin signalling, were shown to modulate EBC proliferation during wound healing (Elbediwy et al., 2016). However, their role in early epidermal development is poorly characterised.

In addition to their role in cell proliferation, YAP1 and WWTR1 promote cell survival. When cells are challenged with DNA damaging agents including doxorubicin (Ma et al., 2016) and radiation (Guillermin et al., 2021), YAP1, as a co-transcription factor of Teads, is localised in the nucleus to drive expression of pro-survival genes. Indeed, transgenic expression of *Yap1* in murine livers upregulates the expression of *Survivin* (*Birc5*) in hepatocytes, protecting these cells from apoptosis when treated with a pro-apoptotic agent, Jo-2 (Dong et al., 2007). On the other hand, genetic knock-down of *Yap1* or inhibition of YAP1 nuclear localisation exaggerates apoptosis in cells exposed to chemo- and radiotherapy (Guillermin et al., 2021; Ma et al., 2016). Thus, YAP1 executes its pro-survival function in the nucleus by driving the expression of target genes.

In this study, we show that concurrent loss of *yap1* and *wwtr1* results in aberrant apoptosis of EBCs on the yolk at the 6-10 ss. However, epidermal cell proliferation is not reduced in *yap1;wwtr1* double mutant zebrafish embryos, in contrast to adult *Yap1^−/−^;Wwtr1^−/−^* mice. At these developmental stages, Yap1 and Wwtr1 are localised in the EBC nuclei to putatively carry out their co-transcriptional function through the Tead-Binding Domain and support epidermal cell survival. This cell death phenotype in mutant embryos is executed through the extrinsic apoptotic pathway, as apoptosis assays reveal activation of Caspase-8 and Caspase-3. Further, we show that γH2AX, a DNA damage marker, is recruited in mutant EBCs. Thus, DNA damage prompted the extrinsic apoptotic pathway in these cells (Hill et al., 1999). Live imaging of epidermal cells found that, leading up to this apoptosis phenotype, mutant basal cell growth is inhibited after the 6 ss, unlike their sibling embryo counterparts, which continue to grow. Interestingly, human keratinocytes cultured on stiffer substrates exhibit more γH2AX foci than cells cultured on softer substrates, suggesting that the mechanical environment not only modifies cell size, but also plays a role in genomic stress.

Thus, we propose that in response to genomic stress, Yap1 and Wwtr1 have important roles in the survival of EBCs during the course of their development.

## RESULTS

### Yap1 and Wwtr1 are localised in the nuclei of epidermal basal cells to modulate their survival

The previous work on *yap1;wwtr1* double mutant zebrafish reported no overt cell death in the tail (Kimelman et al., 2017). We focused on the lateral yolk, on which two layers of the epidermis – the periderm and basal epidermis – reside. We performed TUNEL assays on 16-18 ss embryos and found aberrant apoptosis in the *yap1^−/−^;wwtr1^−/−^* (henceforth referred simply as ‘mutant’) epidermis on the yolk, but not in the tail as reported before (Figure 1A). Additionally, whereas WT embryos have regular distribution of P63-positive nuclei (an EBC marker) over the yolk, mutants have a more irregular and sparser pattern (Figure 1A), suggesting apoptosis of mutant EBCs. We investigated this phenotype in closer detail by repeating this assay in younger embryos and imaging them with higher magnifications. Normal sibling epidermis exhibit virtually no TUNEL signal. On the other hand, mutant epidermis exhibit P63 and TUNEL co-staining (apoptotic EBCs), as well as punctate TUNEL signal. The latter corresponds to apoptotic fragments that made up the bulk of the TUNEL signal (Figure 1B and B’). Therefore, Yap1 and Wwtr1 are required for EBC survival.

**Figure 1.**
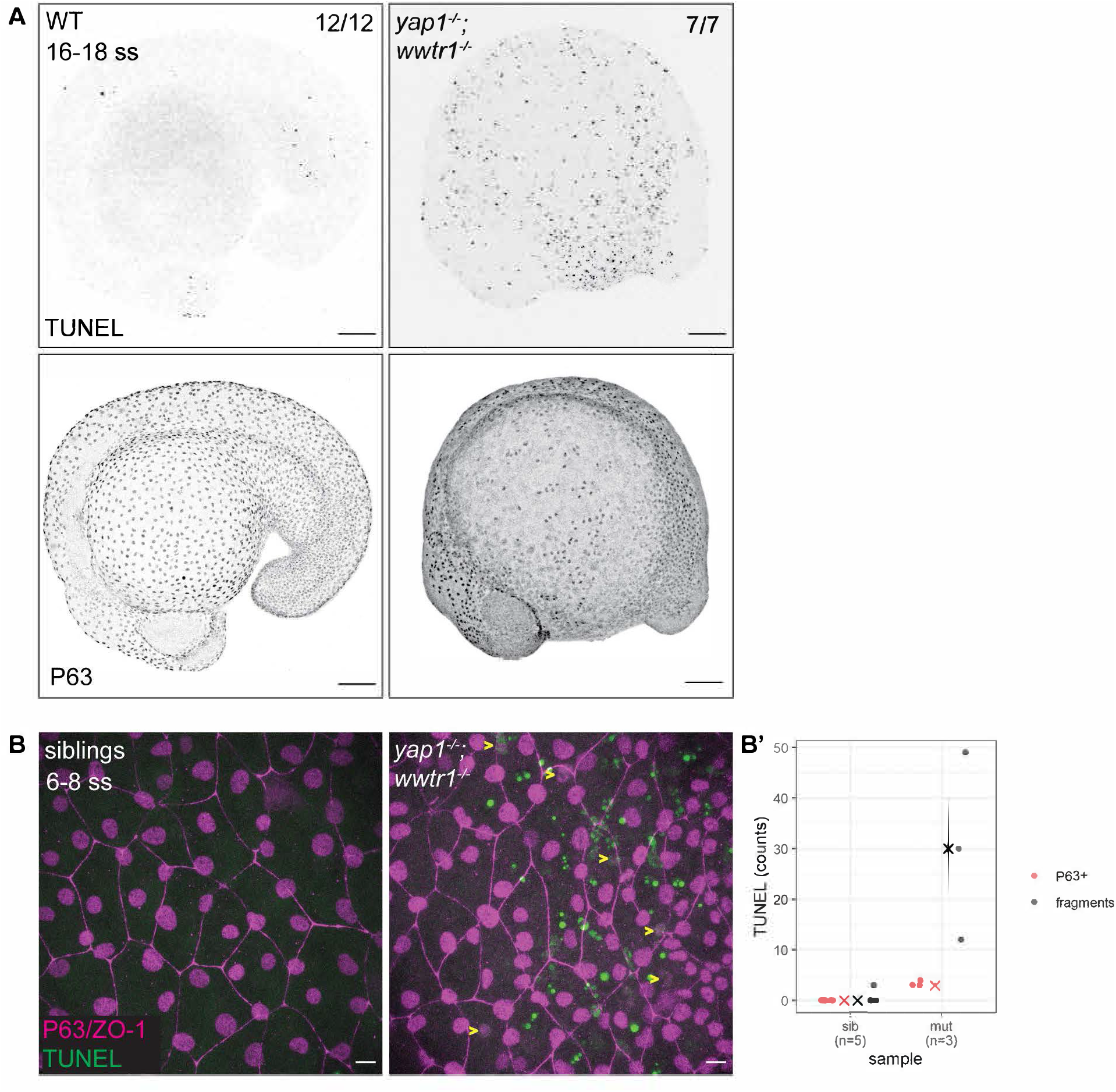
Developing zebrafish *yap1;wwtr1* double mutants show aberrant epidermal basal cell death. (A) Maximum intensity projections of 16-18 ss WT and mutant embryos stained with TUNEL and P63. Scale bars, 100 μm. (B) Maximum intensity projections of the epidermis on the lateral yolk of 6-8 ss mutant and normal siblings stained with TUNEL, P63 and ZO-1. Yellow arrowheads indicate TUNEL and P63 double positive nuclei. Scale bars, 10 μm. (B’) Quantification of TUNEL-positive EBC nuclei (P63+) and apoptotic fragments (fragments).

Using previously characterised antibodies against Yap1 and Wwtr1 (Flinn et al., 2020; Kimelman et al., 2017; Lai et al., 2018; Miesfeld et al., 2015), we investigated their cellular localisation in the epidermis. Both Yap1 and Wwtr1 are localised in the nuclei of the peridermis and basal epidermis at the tailbud stage and at the 16-20 ss (Figure 2). Yap1 is also localised to the junctions of peridermal cells (Figure 2; Flinn et al., 2020). At these developmental stages, some EBCs in the basal epidermis lose P63 expression and differentiate into ionocytes (Jänicke et al., 2007). By comparing P63-positive to P63-negative nuclei in the basal epidermis, we found greater expression levels of Yap1 and Wwtr1 in the P63-positive ones (Figure 2-figure supplement 1A), indicating Yap1 and Wwtr1 activity in EBCs.

**Figure 2.**
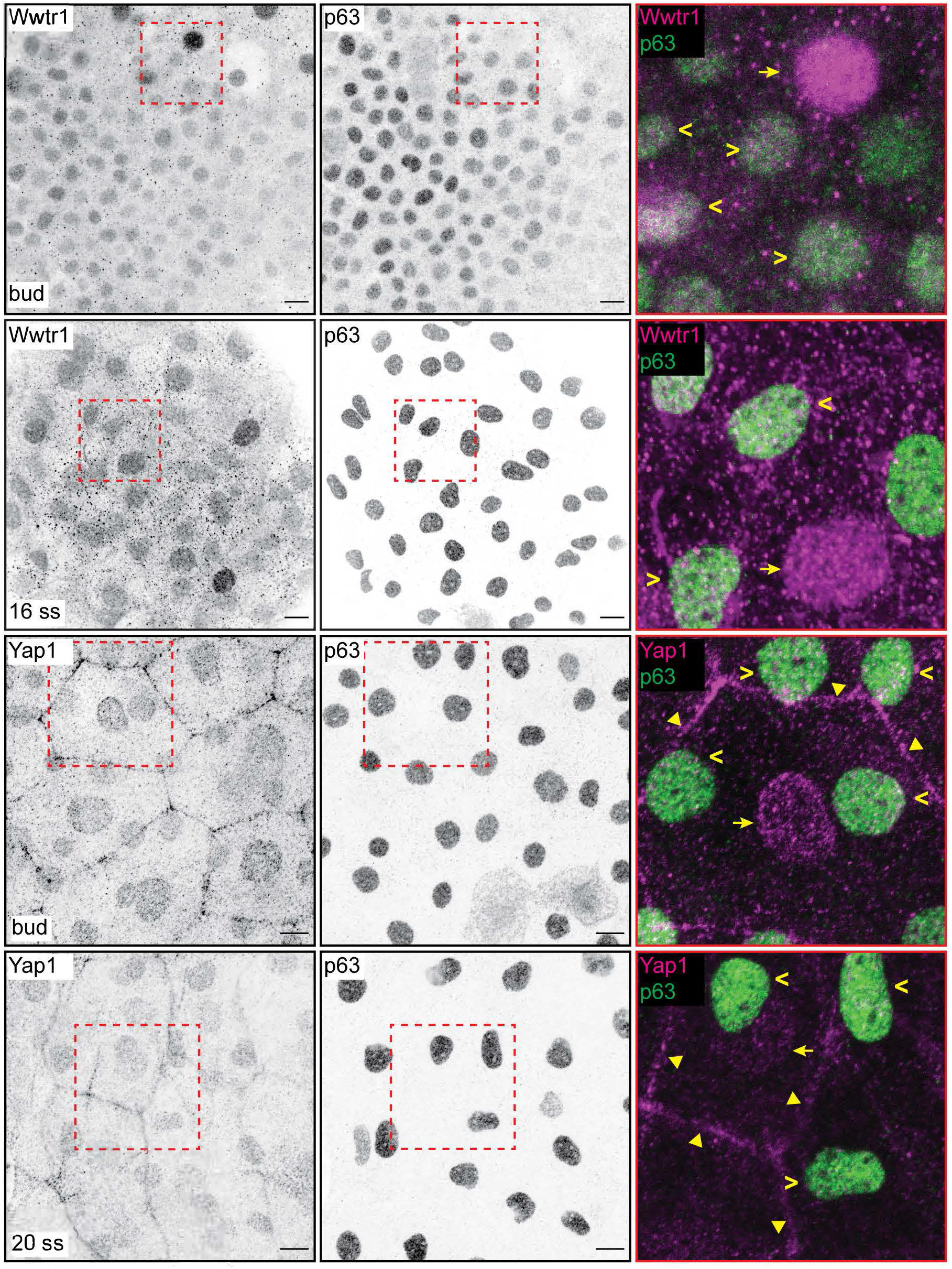
Yap1 and Wwtr1 are localised in EBCs and EVL cells. Maximum intensity projections of the epidermis on the lateral yolk stained with P63, and Wwtr1 or Yap1 antibodies at indicated developmental stages. Insets, demarcated in red, show overlay of P63 and Yap1/Wwtr1. Arrowheads – P63-positive basal cells; arrows – peridermal cells; triangles – peridermal cell junctions. Scale bars, 10 μm.

In the nucleus, Yap1 and Wwtr1 bind to Teads via their Tead-Binding Domain (TBD) to drive expression of target genes. To test whether this interaction is important for their function, we evaluated the *yap1^bns22^* zebrafish allele which encodes an in-frame deletion in the TBD of Yap1 (p.(Pro42_Glu60delinsLeu)) (Figure 2-figure supplement 1B). Specifically, this deletion spans across the S54 residue that binds to Teads (Miesfeld et al., 2015; Zhao et al., 2008). In the *yap1^bns22/bns22^;wwtr1^−/−^* embryos, we again observed apoptosis in the epidermis (Figure 2-figure supplement 1B’). These observations suggest that nuclear-localised Yap1 in EBCs promote survival by driving the expression of downstream target genes with Teads.

### Yap1 and Wwtr1 are dispensable for the proliferation of developing epidermal cells

Using adult mice, a previous study found fewer proliferating cells in the epidermis of *Yap1;Wwtr1* double mutants (Elbediwy et al., 2016). Thus, we performed a proliferation assay with phospho-histone H3 (pH3) staining in the 3-5 ss embryos. We chose this developmental stage to avoid coinciding with the cell death phenotype in mutants. Overall, the number of pH3 foci is not noticeably different between WT and mutant embryos (Figure 3). Restricting our analysis to the epidermis on the lateral yolk, the number of pH3 positive cells is not significantly different between the two genotypes (Figure 3B). Thus, Yap1 and Wwtr1 are not required for epidermal cell proliferation during development.

**Figure 3.**
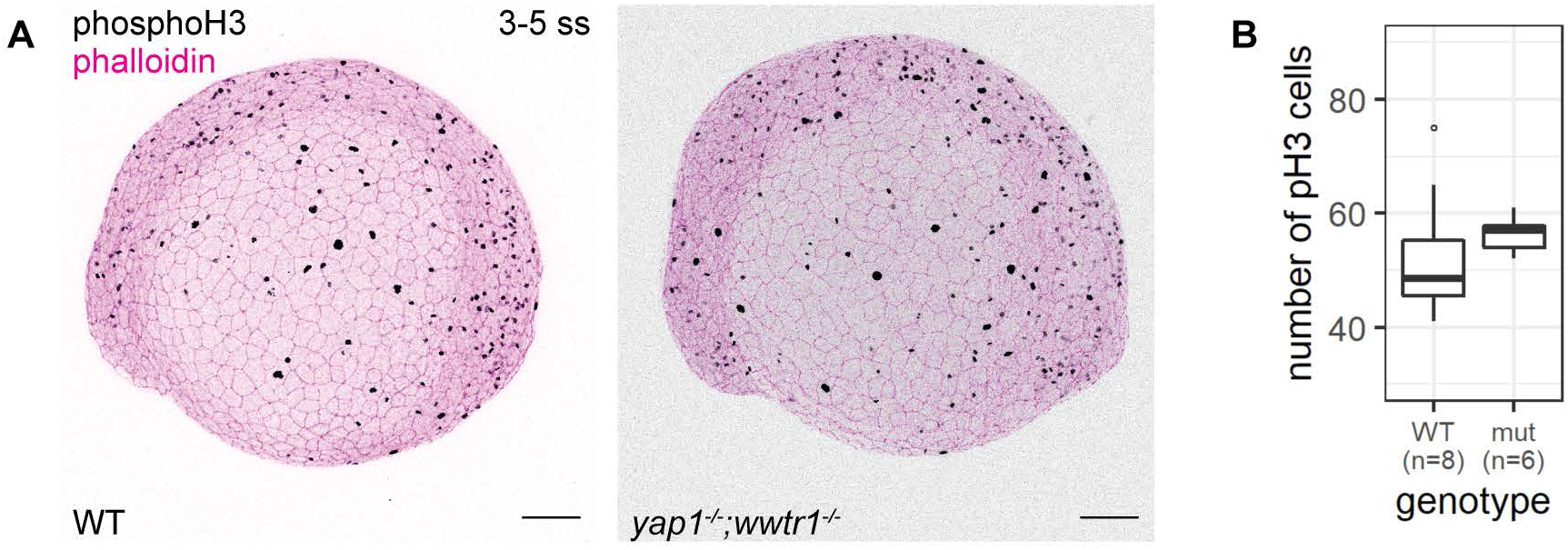
Concurrent loss of *yap1* and *wwtr1* did not impair epidermal cell proliferation. (A) Maximum intensity projections of 3-5 ss WT and mutant embryos stained with a proliferation marker, phospho-histone H3 (phosphoH3; pH3), and phalloidin. Scale bars, 100 μm. (B) Boxplot of the number of pH3 cells on the lateral side of the yolk of WT and mutant zebrafish embryos. mut – *yap1;wwtr1* double mutants.

### Apoptosis in mutant EBCs occurs through the extrinsic pathway

We next address whether the apoptosis phenotype in mutant EBCs is initiated by the intrinsic or extrinsic pathway. Supporting the TUNEL results, we find in mutant epidermis, cleaved Caspase-3 co-stained with P63 and in apoptotic fragments (Figure 4A and A’). To test the extrinsic pathway, we evaluated the activity of Caspase-8 using a live Caspase-8 probe, FITC-IETD-FMK. At the 18-20 ss, FITC foci are present in mutant, but not WT, epidermal cells (Figure 4B and B’), indicating that Caspase-8 is active in mutants. In addition, the apoptosis phenotype is indistinguishable between mutant embryos with and without morpholino knock-down of *tp53* (Figure 4-figure supplement 1A). Taken together, apoptosis in *yap1;wwtr1* double mutant epidermal cells is initiated by the extrinsic pathway and is independent of Tp53.

**Figure 4.**
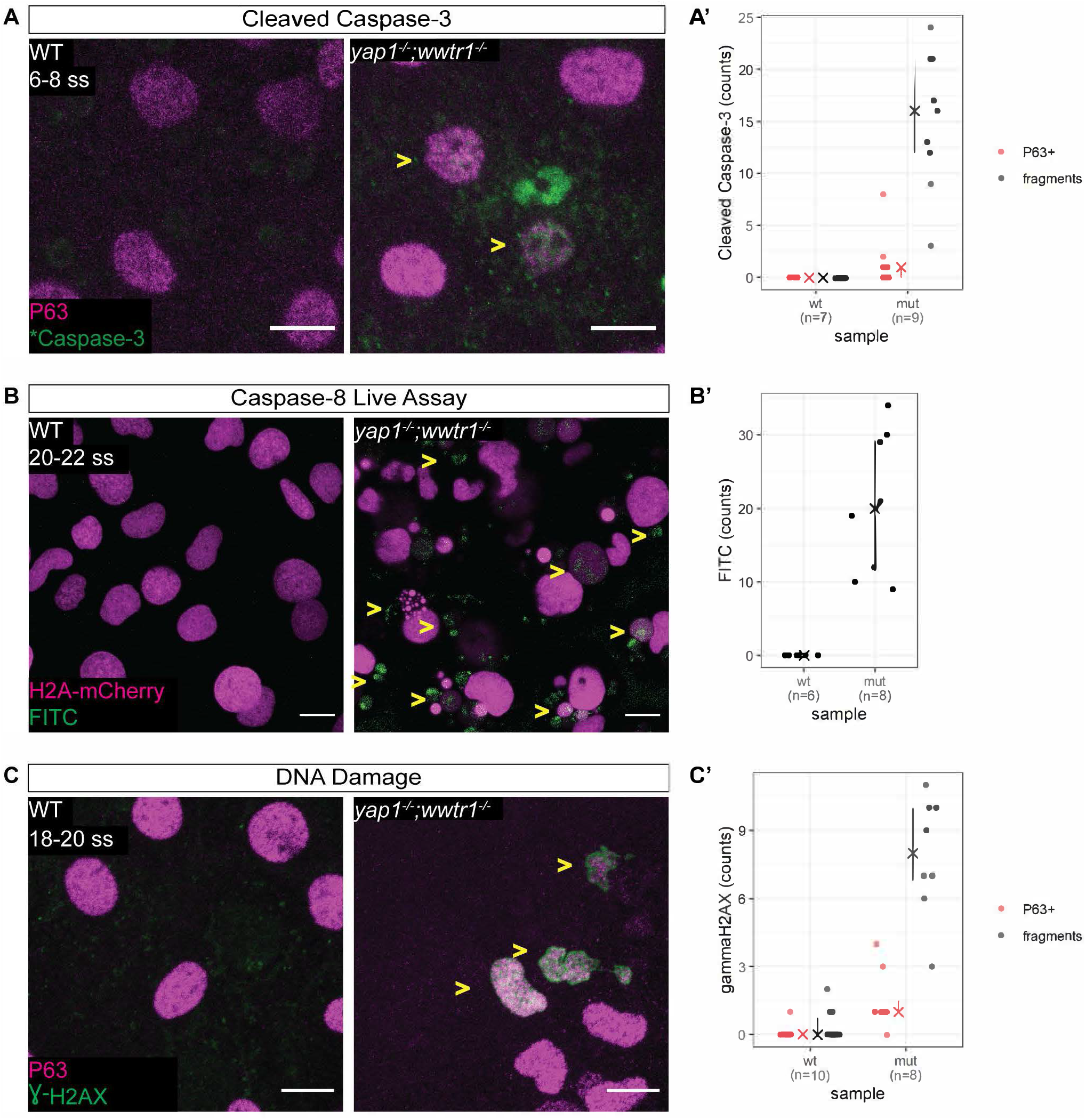
Zebrafish *yap1^−/−^;wwtr1^−/−^* epidermal cells exhibit DNA damage and extrinsic apoptotic cell death. (A) Maximum intensity projections of 6-8 ss WT and mutant epidermis on the lateral yolk stained with cleaved Caspase-3 (*Caspase-3) and P63. Some *Caspase-3 positive cells are also P63 positive (yellow arrow heads). Scale bars, 10 μm. (A’) Number of EBCs (P63+) and apoptotic fragments (fragments) expressing cleaved Caspase-3 in WT and mutant embryo epidermis. ‘X’ represents the median, while the whiskers projecting from it represent the interquartile range. (B) Maximum intensity projections of the epidermis on the lateral yolk of 20-22 ss WT and mutant embryos expressing H2A-mCherry. Embryos were incubated in a Caspase-8 chemical probe, FITC-IETD-FMK, prior to imaging. FITC signal indicates Caspase-8 activity in cells (yellow arrowheads). Scale bars, 10 μm. (B’) Number of FITC foci in WT and mutant embryo epidermis. ‘X’ represents the median, while the whiskers projecting from it represent the interquartile range. (C) Maximum intensity projections of 18-22 ss WT and mutant epidermis on the lateral yolk stained with γH2AX and P63. Some nuclei are positive for both markers (yellow arrowheads). Scale bars, 10 μm. (C’) Number of EBCs (P63+) and apoptotic fragments (fragments) expressing γH2AX in WT and mutant embryo epidermis. ‘X’ represents the median, while the whiskers projecting from it represent the interquartile range.

### γH2AX is recruited to the nuclei of mutant epidermal cells

Previous studies have found that the epidermis clears DNA-damaged cells through the extrinsic apoptotic pathway (Bang et al., 2003; Hill et al., 1999; Takahashi, 2001), prompting us to investigate our mutant epidermal cells for DNA damage. Indeed, interrogating mutant and WT embryos by Western Blot shows elevated levels of γH2AX, a DNA damage marker, in mutants (Figure 4-figure supplement 1B). Closer inspection of these embryos by confocal microscopy reveals a pan-nuclear pattern of γH2AX in mutant EBCs but not in WT ones (Figure 4C and C’). γH2AX was also observed in apoptotic fragments. Therefore, DNA damage is the putative stimulus of the extrinsic apoptotic pathway in mutant EBCs.

### Mutant basal cell size growth is limited

What are the cellular events leading to apoptosis in mutant embryos? We utilised live imaging of the developing epidermis in order to visualise the cell behaviour. Embryos at the one-cell stage were injected with *H2A-mCherry* and *lyn-EGFP* mRNA to mark the nuclei and membrane, respectively. At the tailbud stage, we mounted the embryos laterally on the yolk to record the epidermal cells. In sibling and *yap1;wwtr1* double mutants, epidermal cell behaviour is not readily distinguishable between them throughout the live imaging experiment, aside from apoptosis in mutant embryos beginning from the 3-hour mark (Movie S1).

With the membrane marker, we measured basal cell size in sibling and mutant embryos. Whereas basal cells in sibling embryos grew steadily throughout the live-imaging experiment, mutant ones stopped growing 120 minutes post-tailbud (Figure 5). At the end of this experiment, basal cells in sibling embryos were on average a third larger than mutant ones (747 ± 136 μm^2^ vs. 551 ± 103 μm^2^ [mean ± s.d.]; *P* < 0.01). Curiously, apoptosis – DNA condensation and membrane blebbing (Movie S2) – of mutant basal cells follows after stagnation of their growth (Figure 5). We investigated cell shape parameters as well, but we did not observe any clear differences between mutant and sibling embryo basal cells. Taken together, basal cell growth is inhibited in *yap1;wwtr1* double mutant embryos.

**Figure 5.**
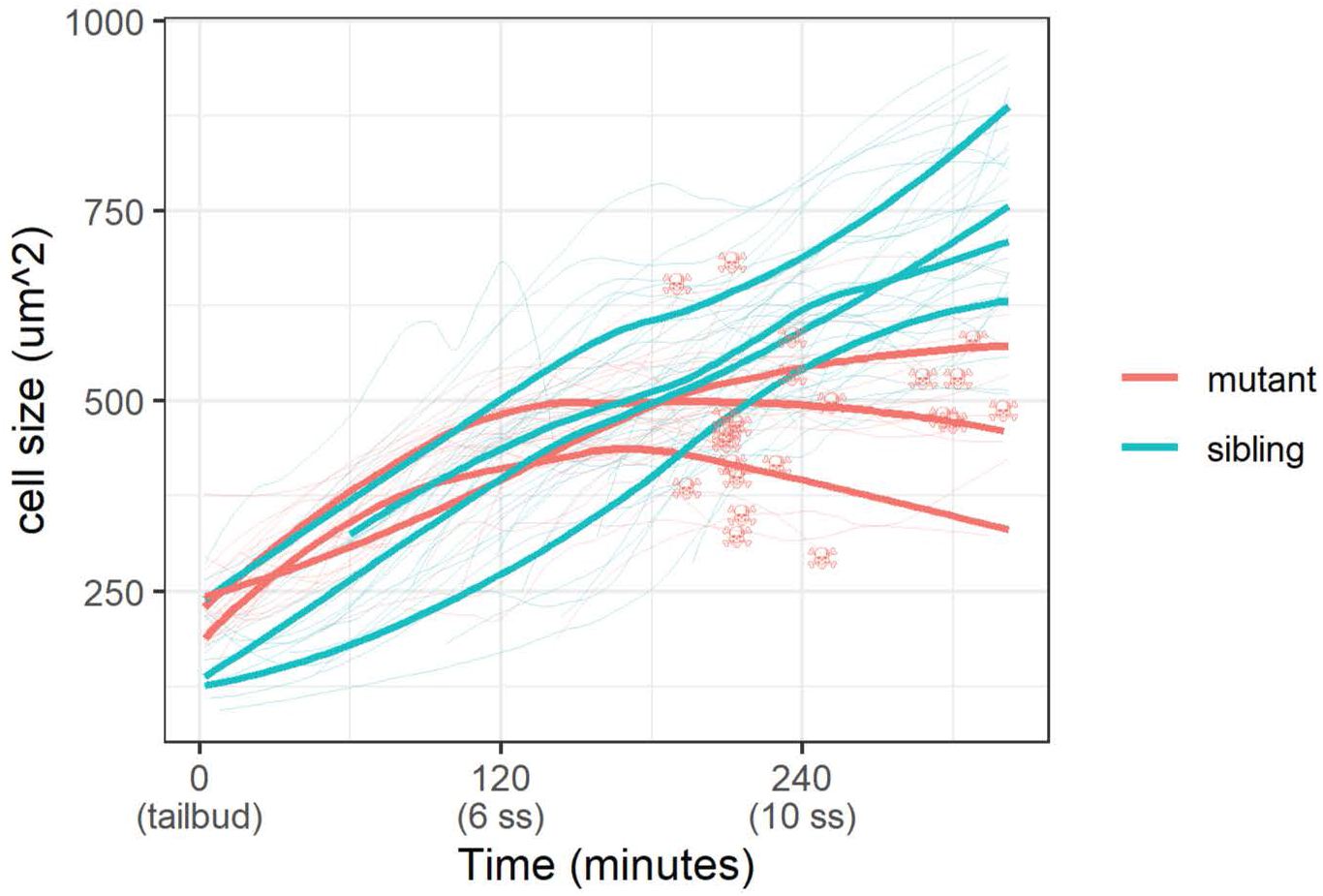
Epidermal cell size of sibling and mutant embryos during development. Cell size was measured with a membrane marker during live imaging of mutant and sibling embryos from the tailbud stage. Fine lines are the size of individual cells, while bold lines are the average cell size in a single embryo. Skull symbols mark cell size and time before death (DNA condensation).

### Investigation of γH2AX by cell stretching and substrate stiffness

Our results show that DNA damage is an upstream trigger of extrinsic apoptosis in mutant EBCs, and that apoptosis occurs after inhibition of cell size. Thus, we hypothesised that factors underlying cell size play a role in DNA damage. We tested this hypothesis with two experiments: (1) *in vivo* stretching of head epidermal cells by ventricular inflation; and (2) culturing of a human keratinocyte cell line, HaCaT, on increasing stiffness of collagen-coated hydrogels.

We stretched epidermal cells *in vivo* by inflating the zebrafish brain ventricle with mineral oil (Figure 6A) (Lewis et al., 2020). As a positive control, we exposed zebrafish embryos to UV for 10 minutes followed by recovery of 1 hour before fixation. In the UV-treated embryos, we found pan-nuclear staining of γH2AX in the periderm and basal epidermis (Figure 6-figure supplement 1A), similar to the staining pattern in mutant EBCs (Figure 4C). However, we did not observe γH2AX in the nuclei of epidermal cells after ventricular inflation (Figure 6A’). These observations suggest that stretching of epidermal cells alone does not induce the recruitment of γH2AX.

**Figure 6.**
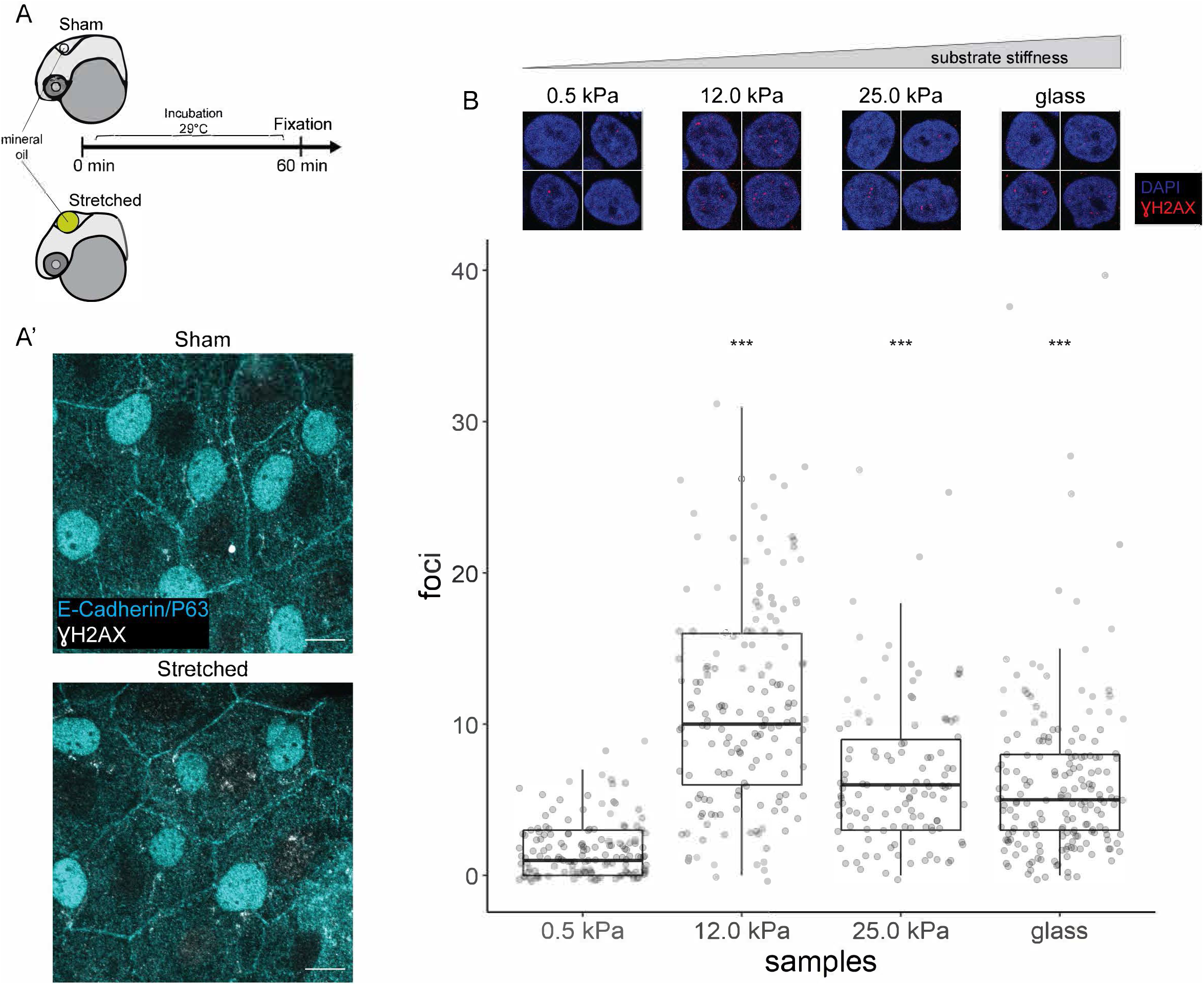
γH2AX in stretched epidermal cells and in keratinocytes cultured on different substrate stiffness. (A) Schematic of head epidermal cell stretching experiment in zebrafish embryos. (A’) Sham (n = 6) and stretched (n = 6) head epidermal cells stained with γH2AX and epidermal markers (E-Cadherin and P63). P63 is a marker for EBCs. No discernible γH2AX signal was detected in both conditions. Scale bars, 10 μm. (B) Selected nuclei of HaCaT cells cultured on 0.5 kPa, 12.0 kPa, 25.0 kPa hydrogels and glass, as well as boxplot of the number of γH2AX foci in these nuclei. Number of γH2AX foci in HaCaT cell nuclei were contrasted against HaCaT cells cultured on 0.5 kPa hydrogel using T-tests. Data were collected from 3 independent experiments. *** - *P* < 0.001 adjusted for multiple testing (Tukey method).

Next, we evaluated DNA damage in HaCaT cells cultured on increasing substrate stiffness - 0.5 kPa, 12 kPa, 25 kPa hydrogels and glass. One set of cells was exposed to UV as positive controls. Interestingly, the number of γH2AX foci detected in HaCaT cell nuclei is higher in cells cultured on stiffer substrates, but this correlation is not linear, with the maximum number of foci occurring in HaCaT cells cultured on 12 kPa hydrogel (Figure 6B). Moreover, we observed a similar trend among cells exposed to UV (Figure 6-figure supplement 1B). Nevertheless, in cells cultured on a given substrate stiffness, we detected more γH2AX foci in the UV group than the unexposed group (Figure 6-figure supplement 1B). Thus, keratinocytes cultured on stiff substrates appear to have greater genomic stress than cells on soft substrates.

## DISCUSSION

We report that Yap1 and Wwtr1 play important roles in epidermal cell size control and survival during epidermal development. In contrast to adult murine epidermis (Elbediwy et al., 2016), the developing zebrafish epidermis does not require Yap1 and Wwtr1 for cell proliferation. Instead, we found mutant epidermal cells to be smaller, exhibit DNA damage and undergo apoptosis through the extrinsic pathway. These observations led us to investigate the relationship between the mechanical environment of epidermal cells and genomic stress.

The mechanical environment of the developing epidermis changes with increasing production of the extracellular matrix (ECM). Indeed, greater ECM density is known to increase cell size (Engler et al., 2006), which could underscore the increasing basal cell size during epidermal development. In the transcriptomic signatures from the previous study (Kimelman et al., 2017), genes that are downregulated in *yap1;wwtr1* double mutant tailbuds include *lama5*, *col7a1*, *lamb1a*, *lamb4* and *lamb2*, which encode key components of the skin basal lamina. This downregulation likely translates to a different ECM environment in mutant embryos and consequently results in the inhibition of basal cell size (Figure 5).

Mechanical perturbations of cells were shown to recruit DNA damage factors including γH2AX and ATR (Cognart et al., 2020; Kumar et al., 2014). Our present results show that both baseline and induced DNA damage can be intensified by culturing cells on stiffer substrates. Specifically, we found more γH2AX foci in human keratinocyte cells seeded on 12 kPa, 25 kPa hydrogels and glass compared to the ones seeded on 0.5 kPa hydrogels. Correspondingly, expression of genes encoding components of the skin basal lamina increases during the early segmentation stages (Baseline expression from transcriptional profiling of zebrafish developmental stages, Expression Atlas) (Papatheodorou et al., 2020). This increase may create an increasingly rigid environment (Engler et al., 2006) that perhaps results in greater baseline genomic stress in the developing EBCs, and thus require survival signals to counteract this stress. Previous investigations have already shown how Yap1 plays instrumental roles in providing this survival signal, including driving the expression of *Survivin*, under genomic stress (Guillermin et al., 2021; Ma et al., 2016). However, our Western Blot assay did not find downregulation of Survivin in zebrafish *yap1;wwtr1* double mutants (data not shown), presumably as Survivin is also a key player in cellular proliferation (Nair et al., 2013), which is not impeded in mutants. Therefore, we propose that the mechanical environment of the epidermis modifies genomic stress in cells, and that Yap1 and Wwtr1 play a role in their survival.

Our work reveals that Yap1 and Wwtr1 have important functions in epidermal cell survival and growth during development. We also show greater genomic stress in cells grown on more rigid extracellular substrates, which likely explains the DNA damage-induced apoptosis phenotype in *yap1;wwtr1* double mutant zebrafish. These results open new areas of exploration on the physiological relevance of mechanobiology in development, physiology and disease, as well as its interplay with Yap1/Wwtr1 and genomic stress in these processes.

## MATERIALS & METHODS

### Ethics statement

All zebrafish husbandry was performed under standard conditions in accordance with institutional (Biological Resource Center, A*Star, Singapore, and Tata Institute of Fundamental Research, India) and national ethical and animal welfare guidelines.

### Zebrafish lines

Mutant zebrafish lines used in this study are: *yap1^bns19^* (Astone et al., 2018; Kimelman et al., 2017), *yap1^bns22^* (this study) and *wwtr1^bns35^* (Lai et al., 2018).

The *yap1^bns22^* allele was generated from a CRISPR/CAS9-mediated mutagenesis. Two single-guide RNA (sgRNA) sequences, 5’-ACCTCATCGGCACGGAAGGG-3’ and 5’-CTGGAGTGGGACTTTGGCTC-3’, were designed by a sgRNA design tool by the Zhang Lab (crispr.mit.edu; accessed in 2014). These sequences were cloned into the pT7-gRNA vector (Addgene #46759). Following vector linearisation with BsmBI enzyme, sgRNA were obtained by *in vitro* transcription with MEGAshortscript T7 kit (Ambion). *CAS9* mRNA was obtained by *in vitro* transcription of linearised pT3TS-nCas9n vector (Addgene #46757) with MEGAscript T3 kit (Ambion). sgRNA (50 pg each) and 150 pg of *CAS9* mRNA were co-injected into one-cell stage AB embryos. The *yap1^bns22^* allele contains a 54 bp deletion in the Tead-Binding Domain [p.(Pro42_Glu60delinsLeu)]. This allele can be genotyped by using the following PCR primer pair: 5’-CTGTTTGTGGTTTCTGAGGGG-3’ and 5’-CGCTGTGATGAACCCGAAAA-3’. Mutant and WT products can be resolved and distinguished by gel electrophoresis with a 2% gel. Genotyping of *yap1^bns19^* and *wwtr1^bns35^* alleles have been described previously (Astone et al., 2018; Lai et al., 2018).

Mutant embryos in this study were obtained from an incross of double heterozygous animals (i.e. *yap1^bns19/+^*;*wwtr1^bns35/+^* incross or *yap1^bns22/+^*;*wwtr1^bns35/+^* incross).

### Incubation and visual identification of mutant embryos

Experiments involving 16 ss and older embryos were incubated at 25°C throughout, otherwise younger embryos were incubated at 28.5°C.

*yap1;wwtr1* double mutant embryos older than 16 ss can be visually identified for experiments. These mutant embryos were compared to stage-matched embryos from WT crosses.

For experiments involving < 16 ss embryos, embryos from heterozygous incross were fixed and assayed for cell death with TUNEL. Using a wide-field stereomicroscope, embryos with excess TUNEL staining in the epidermis on the lateral yolk were selected and mounted for confocal imaging. Under the confocal microscope, embryos with apoptotic bodies (TUNEL foci and fragmented DNA stained by DAPI) were further investigated and genotyped for downstream analyses. These embryos were compared to stage-matched WT embryos.

### Morpholinos

Morpholinos used in this study were: *tp53* (Robu et al., 2007) and *lama5* (Nagendran et al., 2015; Webb et al., 2007). 1 ng of each morpholino was injected into one-cell stage embryos.

### Whole-mount Immunofluorescence Assay

Zebrafish samples were fixed in 4% PFA in 1X PBS, followed by dechorionation. Briefly, samples were blocked for 1 hour in 2 mg ml^−1^ BSA and 1% Goat Serum (Thermo Fisher) in 1X PBSTXD (1X PBS, 0.1% Tween-20, 0.1% Triton X-100 and 1% DMSO). Then, samples were incubated in primary antibody (see below) overnight at 4°C. Samples were washed in 1X PBSTXD every 15 minutes for 2 hours and then incubated in secondary antibody in room temperature for at least 4 hours, followed by washing with 1X PBSTXD. Counter stains for nuclei was DAPI, and for Actin was Phalloidin (Alexa488 conjugated). When using anti-γH2AX as primary antibody, all PBS buffers were substituted with TBS buffers, and blocking buffer consisted of 3-5% BSA only.

TUNEL assays were performed between the blocking step and primary antibody incubation step. Samples were washed in 1X PBST (1X PBS, 0.1% Tween-20) three times, 5 minutes each. After aspiration of the buffer, samples were incubated in the Fluorescein TUNEL kit (Roche) for 1 hour at 37°C with gentle shaking. 1X PBST washes were performed to stop the reaction before proceeding with the remaining step of the immunofluorescence protocol.

Primary antibodies used in this study were: anti-Yap1 (CS4912, Cell Signalling), 1:250; anti-Wwtr1 (D24E4, Cell Signalling), 1:250; anti-P63 (ab735, Abcam), 1:500; anti-ZO1 (1A12, Thermo Fisher), 1:250; anti-phosphoH3 (9701, Cell Signalling), 1:500; anti-cleaved-Caspase-3 (ab13847, Abcam), 1:250; and anti-γH2AX (GTX127342, Genetex), 1:500, anti E-cadherin (610182, BD Transduction Labs), 1:100.

Secondary antibodies used in this study were: Alexa fluorophore-conjugated anti-mouse and anti-rabbit by Thermo Fisher used at 1:500 dilution.

### Caspase-8 Assay

Embryos were injected with 10-15 pg *H2A-mCherry* mRNA at the one-cell stage. At 16 ss, mutant and WT embryos were identified and dechorionated, then incubated in FITC-IETD-FMK (ab65614, Abcam) for 30 minutes at room temperature. Embryos were washed with egg water 3 times, 5 minutes each, and then mounted for confocal imaging.

### Proliferation assay

Embryos were imaged from the lateral side. After imaging, embryos were genotyped and only *yap1;wwtr1* double mutants were compared with stage-matched WT embryos. A maximum intensity projection was generated for each embryo. The yolk was manually demarcated as a ‘region of interest’, followed by manual identification and counting of phospho-HistoneH3 foci. Poisson regression for count data was used to test whether the number of phospho-HistoneH3 foci differ between WT and mutants.

### Yap1/Wwtr1 immunostaining quantification

Nuclei were segmented with DAPI in order to get total intensity of Yap1, Wwtr1 and P63. peridermal nuclei were excluded from the analysis. The intensity of interests are normalised with DAPI intensity. Nuclei were grouped as P63-positive and P63-negative. T-tests were performed between the P63-positive and P63-negative groups for each normalised Yap1 and Wwtr1 intensity.

### Confocal imaging

Live and fixed embryos were embedded in 1% LMA (Sigma) on glass-bottomed dish (inverted microscopes) or petri dish (upright microscopes). For low-magnification images of the entire embryo, a 10x objective lens was used. Otherwise, high-magnification images were taken with 40x objective (water dipping) or 63x objective (water immersion) lens with at least 1 NA was used. For the ventricular inflation experiment, the stretched region of the head epidermis was flattened and mounted using a coverslip, and imaged on the Zeiss LSM 880 confocal microscope with a Plan-APOCHROMAT 63x/1.4 oil objective at a 1.5x optical zoom.

### Live imaging

Embryos from incrosses of *yap1;wwtr1* double heterozygous animals were injected with 10 pg of *H2A-mCherry* and *lyn-EGFP* mRNA each at the one-cell stage. At 90% epiboly, 6-10 embryos were dechorionated and embedded laterally in 0.5% LMA on a glass-bottomed dish to be imaged under the spinning disk system (Yokogawa CSU-W1). A z-stack of 41 slices, 0.5 μm thick, were acquired per embryo every 2 minutes overnight in an environmental chamber set at 28.5°C. At most 8 embryos were imaged in a single sitting. Embryos were genotyped the next day. All mutants are *yap1^−/−^;wwtr1^−/−^*, while siblings are not *yap1^−/−^* or *wwtr1^−/−^*.

### Cell size quantification

Maximum intensity projections of each embryo were generated for each time point. Basal cells were distinguished from EVL cells manually. The lyn-EGFP marks the cell membrane which was manually demarcated to measure cell size every 5 frames. When a cell divides, it marks the start or end of the tracking. For dying cells in mutants, the tracking stops one frame before the DNA condenses.

### Western Blot

16-18 ss mutant embryos (visually identified) and stage-matched WT embryos were dechorionated and collected in separate Eppendorf tubes. A total of 20 embryos from each group were pooled into one tube to constitute a biological replicate. In 1X PBS, embryos were mechanically de-yolked with a pipette tip. Samples were washed in cold 1X PBS thrice, and then 4X Laemmli Sample Buffer (Biorad) with B-mercapthoethanol (Sigma) was added. Samples were vortexed and boiled at 95°C for 5 minutes three times. Samples were stored in −20°C until ready for use.

Proteins from each sample were resolved using a precast 8-16% gradient gel (Biorad), and transferred to a PVDF membrane following manufacturer’s protocol. Membranes were blocked with 3-5% BSA in 1X TBST (1X TBS, 0.5% Tween-20) for an hour, followed by incubation with primary antibody overnight at 4°C. Membranes were washed with 1X TBST before incubation with secondary antibody for at least 1 hour. Membranes were washed again with 1X TBST. ECL substrate (Biorad) was added to the membrane and imaged with the ChemiDoc imaging system (Biorad).

The primary antibodies used in this assay are: anti-B-actin (A5441, Sigma), 1:1000; anti-γH2AX (GTX127342, Genetex), 1:1000.

The secondary antibodies used in this assay are: HRP-conjugated anti-mouse (31430, Thermo Fisher), 1:10000; HRP-conjugated anti-rabbit (31460, Thermo Fisher), 1:10000.

### Ventricular oil injections

The head epidermis was stretched by injecting hydrophobic mineral oil (Bio-Rad; 1632129) into the hindbrain ventricle of 24hpf zebrafish, as previously described (Lewis et al., 2020). In brief, at 24 hpf the dechorionated embryos were anaesthetised in 0.02-0.024% (w/v in E3) tricaine, and vertically mounted in holes punched in a 3% (w/v) agarose plate. This was followed by either stretching the epidermis by injecting 10-12 nL mineral oil in the hindbrain ventricle (Stretched) or injecting only 0.1-0.2 nL oil to act as an injury control (Sham). All embryos used in a set were obtained from the same clutch and were injected using the same capillary needle, to reduce variation. The test and sham embryos were grown in E3 until fixation an hour later.

### HaCaT Cell culture

All cell culture reagents were purchased from Gibco (ThermoFisher Scientific, MA, USA). The human immortalised keratinocyte cell line, HaCaT, was a gift from CT Lim Lab (MBI, Singapore). HaCaT keratinocytes were cultured in complete Dulbecco’s modified Eagle’s medium (DMEM) that was supplemented with 100 I.U./mL aqueous penicillin, 100 μg/ml streptomycin and 10% heat-inactivated fetal bovine serum (FBS). Cells were maintained at 37°C in a humidified atmosphere containing 5% CO2 and harvested with TrypLE (1X). TrypLE was deactivated with complete media. Cells were pelleted by centrifugation at 180 × g for 5 minutes and then re-suspended in complete media. The cell suspensions were used only when their viability, as assessed by trypan blue exclusion, exceeded 95%. Briefly, 0.1 mL of 0.4% of trypan blue solution was added to 0.1 mL of cell suspension. Cell viability and density were calculated with a hemocytometer.

HaCaT cells were then seeded on 0.5 kPa, 12 kPa and 25 kPa collagen-coated Softslips (Matrigen, CA, USA), as well as glass cover slips in individual wells of a 24-well plate. 50,000 cells were seeded on each substrate. Seeded cells were incubated for 10 hours or overnight for cells to completely attach to the substrates.

### HaCaT Cell immunofluorescence assay

HaCaT cells were fixed with 4% PFA in 1X PBS for 15 minutes. After washing off residual PFA with 1X TBSX (1X TBS supplemented with 0.1% Triton X-100), samples were incubated with blocking solution (4% BSA in 1XTBSX) for 1 hour. Samples were then incubated in primary antibody diluted in blocking solution overnight at 4 °C. After washing off the primary antibody solution with 1X TBSX, samples were incubated in secondary antibodies and counterstains diluted in blocking solution for 2 hours, followed by washes with 1X TBSX. Hydrogel-coated coverslips (Softslips) were inverted unto a coverslip for imaging according to manufacturer’s instructions.

### γH2AX foci quantification in HaCaT cells

Nuclei of HaCaT cells were marked as regions of interest for counting the number of γH2AX foci. Background signal was subtracted using a rolling radius of 5 pixels. A focus is at least 5 pixels in area size, and the number of foci was counted for each nucleus. To test the increase of γH2AX foci in the UV-treated group, one-sided T-tests were performed between UV and control groups for each substrate stiffness. To test the change of the number of γH2AX foci by different substrate stiffness, control HaCaT cells cultured on 0.5 kPa hydrogels was used as the reference in two-sided T-tests. *P*-values were adjusted for multiple hypothesis testing with the Tukey method.

### UV Exposure

24 hpf zebrafish embryos or HaCaT cells were exposed to UV in the Biosafety Cabinet. UV dose for zebrafish experiment was 63.2 mJ/cm^2^, while UV dose for HaCaT cells was 30.6 mJ/cm^2^. After UV exposure, samples were allowed to recover for an hour (zebrafish in a 29.0 °C incubator; HaCaT cells in 37.0 °C humidified incubator).

## ACKNOWLEDGEMENTS

The *yap1^bns22^* zebrafish allele was generated by JKHL in Didier Stainier’s lab (MPI, Bad Nauheim), and we thank him for his support. We also thank Boon Chuan Low (MBI, Singapore), Mahendra Sonawane (TIFR, India) and Tom Carney (NTU, Singapore) for insightful discussions and suggestions. This work was funded by the Singapore Ministry of Education Tier 3 grant MOE2016-T3-1-002 and Mechanobiology Institute core funding.

---

**Figure 2-figure supplement 1.**
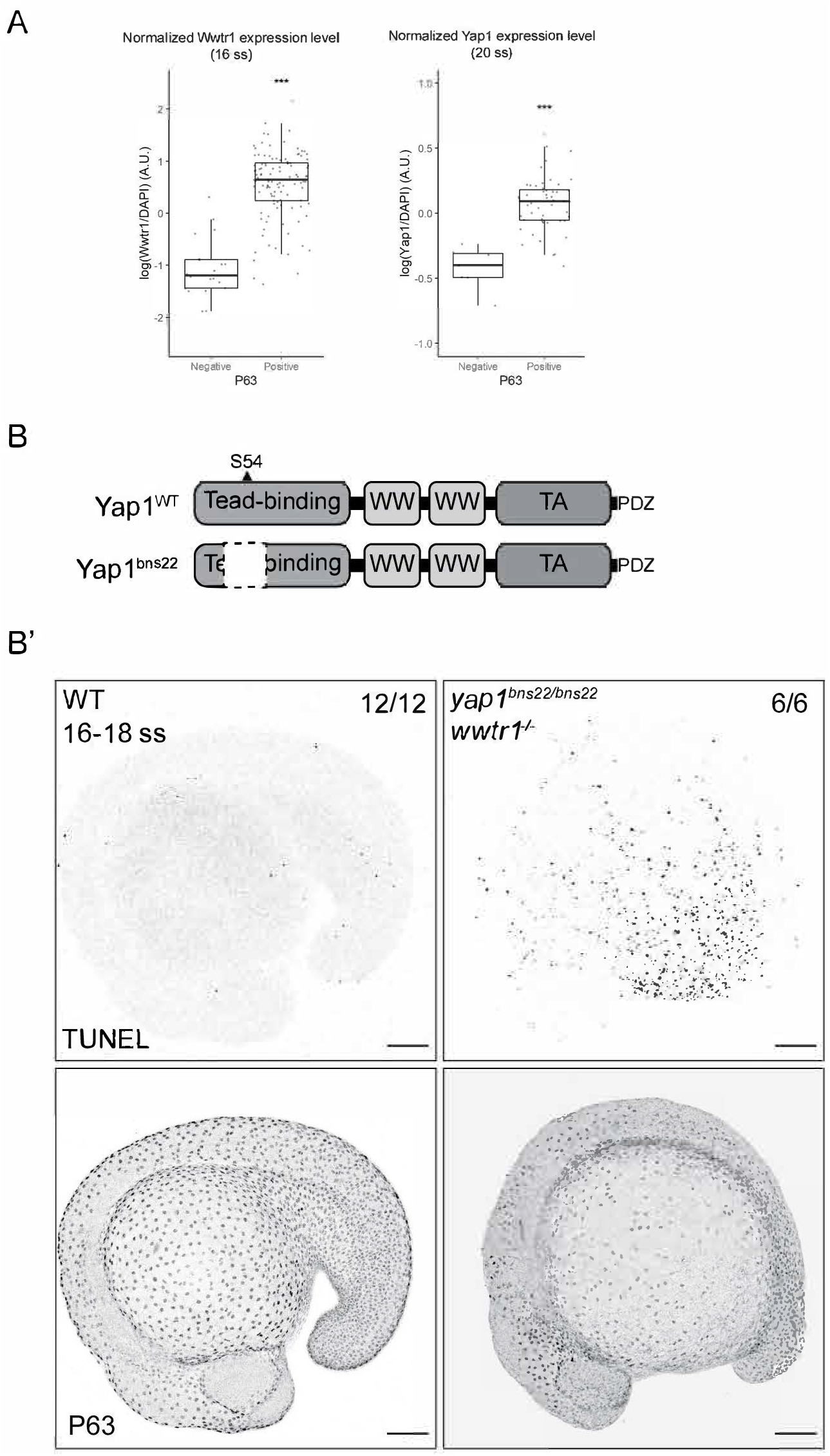
Yap1 and Wwtr1 are localised to the nuclei of EBCs, and Yap1’s Tead-Binding Domain is essential in EBC survival. Boxplots of normalised intensities of Yap1 and Wwtr1 in the nucleus of P63-positive and P63-negative cells in the basal epidermis. T-tests were carried out to compare these intensities between the two groups. *** - *P* < 0.001. (B) Schematic of the encoded protein product of the zebrafish *yap1* WT and bns22 alleles. S54 is necessary for Yap1 binding to Teads. (B’) Maximum intensity projections of 16-18 ss WT and *yap1^bns22/bns22^;wwtr1^−/−^* embryos stained with TUNEL and P63. Scale bars, 100 μm.

**Figure 4-figure supplement 1.**
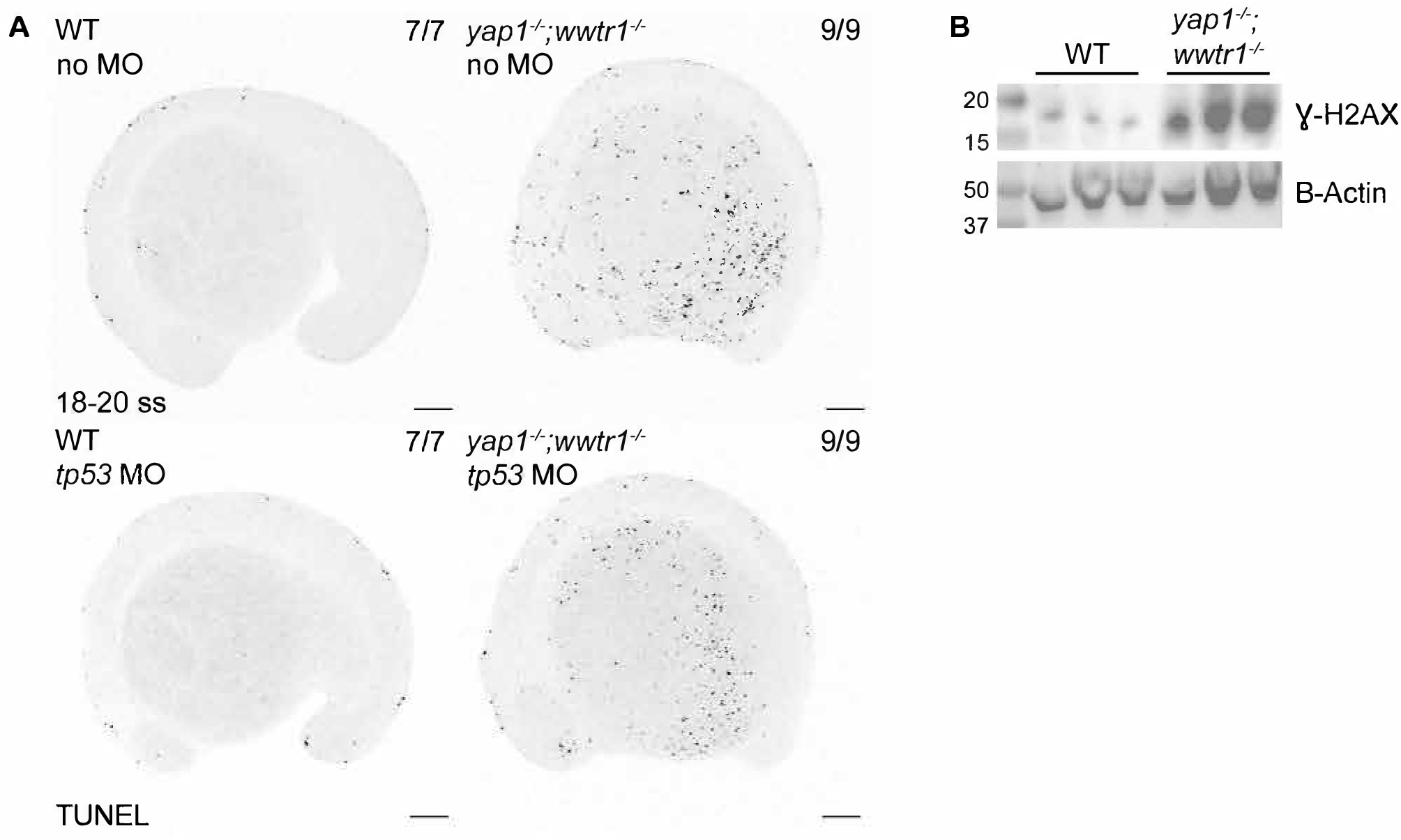
Aberrant cell death and increased γH2AX levels in mutants. (A) Maximum intensity projections of 18-20 ss WT and mutant embryos injected with *H2A-mCherry* mRNA only (no MO) or co-injected with *tp53* morpholino (*tp53* MO). Blocking of *tp53* did not have any observable effect on cell death in both WT and mutant embryos. Scale bars 100 μm. (B) Western Blot of γH2AX and B-Actin (loading control). Each lane is a biological replicate of pooled 18-20 ss WT and mutant embryos. Ladder on far left lane with annotated size (in kD).

**Figure 6-figure supplement 1.**
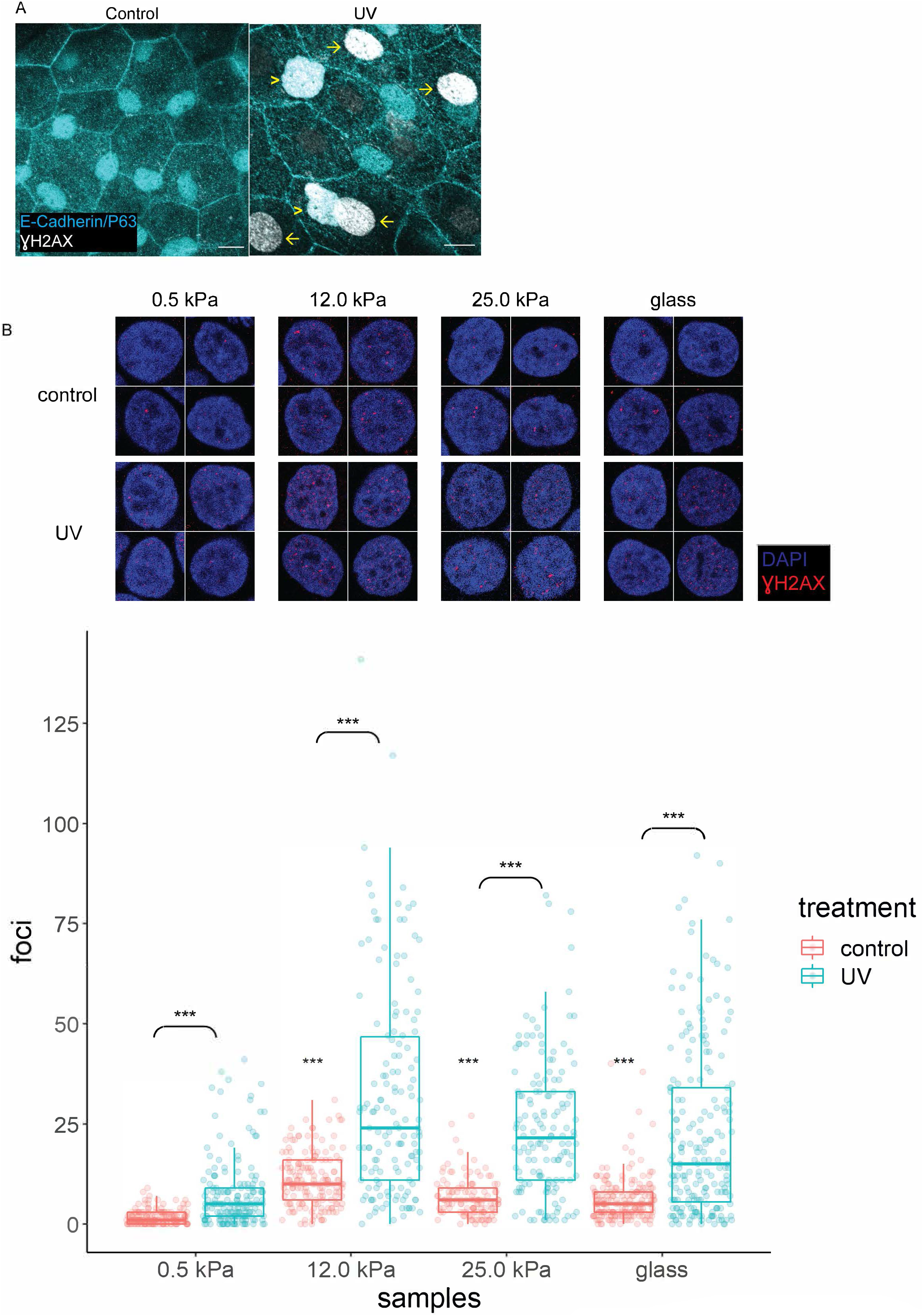
γH2AX in zebrafish head epidermis and HaCaT cells exposed to UV. (A) Control (n = 5) and UV-treated (n = 4) head epidermal cells of zebrafish embryos stained with γH2AX and epidermal markers (E-Cadherin and P63). P63 is a marker for EBCs. EBCs (yellow arrowheads) and peridermal cells (yellow arros) exposed to UV exhibit pan-nuclear γH2AX. (B) Selected nuclei of control and UV-treated HaCaT cells cultured on 0.5 kPa, 12 kPa, 25kPa hydrogels and glass, and boxplots of the number of γH2AX foci in these nuclei. T-tests on control groups were compared against control HaCaT cells cultured on 0.5 kPa hydrogel. T-tests on UV groups were compared against their respective controls (brackets). Data were collected from 3 independent experiments. *** - *P* < 0.001 adjusted for multiple testing.

**Movie S1. Time lapse imaging of mutant and sibling epidermis**. Maximum intensity projections of the epidermis on the lateral yolk of mutant and sibling embryos. Membranes and nuclei are marked by Lyn-EGFP (green) and H2A-mCherry (red), respectively. Time stamp format, HH:MM; 00:00 is tailbud stage.

**Movie S2. Apoptosis of mutant basal cells**. Time lapse of mutant basal cells undergoing apoptosis as captured by live imaging. Membranes and nuclei are marked by Lyn-EGFP (green) and H2A-mCherry (red), respectively. Time stamp format, HH:MM; 00:00 is tailbud stage. Scale bars, 10 μm.

## SOURCE DATA LEGEND

**Figure 1-source data 1**. TUNEL count data for Figure 1B’

**Figure 3-source data 1**. pH3 count data for Figure 3B

**Figure 4-source data 1**. Casp3, Casp8 and γH2AX count data for Figures 4A’, B’ and C’, respectively.

**Figure 5-source data 1**. Measured cell size from live imaging experiment for Figure 5.

**Figure 6-source data 1**. γH2AX foci count data for Figure 6B.

**Figure 2-figure supplement 1-source data 1**. Normalized Yap1 and Wwtr1 intensity measurements for Figure 2-figure supplement 1A.

**Figure 4-figure supplement 1-source data 1**. Raw image files of Western Blot assay for Figure 4-figure supplement 1B.

**Figure 6-figure supplement 1-source data 1**. γH2AX foci count data for Figure 6-figure supplement 1B.

